# LncRNA XIST knockdown suppresses cell proliferation and promotes apoptosis in diabetic cataract via miR-34a/SMAD2 axis

**DOI:** 10.1101/2021.02.01.429101

**Authors:** Chao Wang, Ruiling Zhao, Suhong Zhang

**Affiliations:** Department of Ophthalmology, Shandong Zaozhuang Municipal Hospital, Zaozhuang, P.R. China; Department of Ophthalmology, Shandong Tengzhou Central People’s Hospital, Zaozhuang, P.R. China

**Keywords:** XIST, miR-34a, SMAD2, lens epithelial cells, diabetic cataract

## Abstract

Emerging evidence has manifested that long non-coding RNAs (lncRNAs) played critical roles in diabetes. The present research aimed to investigate the role and mechanism of XIST on proliferation, migration and apoptosis in diabetic cataract (DC). In the present study, lens epithelial cells (SRA01/04) were treated by high glucose (HG). The levels of XIST, miR-34a and SMAD2 were examined by RT-qPCR. MTT, transwell, wound healing and TUNEL assays were employed to examine cell proliferation, invasion, migration and apoptosis. The interaction between miR-34a and XIST or SMAD2 was verified by luciferase reporter assay. It was found that XIST expression was increased and miR-34a level was decreased in DC tissues and HG-induced SRA01/04 cells. XIST knockdown or miR-34a addition attenuated cell proliferation and migration, and induced apoptosis in SRA01/04 cells under HG. XIST targeted miR-34a and regulated DC progression via miR-34a. SMAD2 was a target gene of miR-34a and was positively modulated by XIST. SMAD2 addition accelerated cell proliferation, migration and inhibited the apoptosis in HG-stimulated SRA01/04 cells, which were abrogated by XIST depletion. In conclusion, XIST facilitated the proliferation, migration and invasion, and inhibited the apoptosis via miR-34a/SMAD2 axis in DC.

## Introduction

Diabetes mellitus (DM) causes many complications, including the formation and occurrence of cataracts [1]. Despite the successful surgical replacement of cataracts with intraocular lenses, cataracts are still one of the leading causes of visual impairment and blindness worldwide [2]. Diabetic cataract (DC) characterized by high blood glucose level usually occurs earlier and progresses faster than age-related cataract [3, 4]. Human lens epithelial cells (HLECs) was reported to play essential roles in ocular health and many diseases, including DC [5]. The pathogenesis of DC was a multifactorial process and changed gene expression related to proliferation, differentiation and epithelial-mesenchymal transition (EMT) in LECs may lead to the occurrence of cataract [6]. Thus, it is of importance to explore the mechanism of LECs proliferation in DC.

Long non-coding RNAs (lncRNAs) are a group of non-coding transcripts >200 nts in length that lack protein-encoding potential [7]. Moreover, lncRNAs were reported to participate in DC development. For example, lncRNA MALAT1 facilitated the apoptosis and oxidative stress of HLECs via p38MAPK pathway in DC [8]. SP1-mediated lncRNA PVT1 regulated the viability and apoptosis of LECs in DC through the miR-214-3p/MMP2 axis [9]. Moreover, lncRNA XIST has been found to be associated with diabetic complications, such as diabetic nephropathy and diabetic retinopathy [10, 11]. However, the biological role of XIST during DC is unclear. MicroRNAs (miRNAs) are small RNAs that can target the 3’ UTRs of mRNAs to modulate the transcription of genes [12]. miRNA dysregulation has been uncovered in multiple diseases, including DC. For instance, miR-30a suppressed the autophagy via targeting BECN1 in human DC [13]. MiR-211 promoted apoptosis and repressed proliferation of LECs in DC mice via targeting SIRT1 [14]. Moreover, miR-34a inhibited the viability and induced apoptosis of LECs by targeting E2F3 [15]. Furthermore, SMAD2 was reported to act as a crucial role in posterior capsular opacification [16]. Nevertheless, the regulatory mechanisms of miR-34a and SMAD2 during DC remains largely unknown.

In the present study, we used SRA01/04 cells stimulated by HG as DC model, and the biological effects of XIST on DC was then investigated.

## Materials and methods

### Samples

A total of 32 posterior capsular tissue from diabetic cataract patients and paired normal posterior capsular tissue samples without diabetic cataract were obtained from the Shandong Zaozhuang Municipal Hospital. The tissues were rapidly frozen in liquid nitrogen before used. This study was approved by the Ethics Committee of Shandong Zaozhuang Municipal Hospital and each participator had signed informed consent before the surgery.

### Cell culture and HG treatment

Human lens epithelial cells (SRA01/04) were purchased from Beina Chuanglian Biotechnology Co., Ltd (Suzhou, China). The cells were incubated at 37°C with 5% CO2 in DMEM (Gibco, Waltham, MA, USA) containing 10% FBS (Gibco). To establish DC cell model, SRA01/04 cells were maintained in medium containing high glucose (HG, 25 mM) for 24 h, and cells in normal glucose condition (NG, 5.5 mM) were acted as controls.

### Cell transfection

shRNA against XIST (shXIST; 5’-GUGCGUACAGUGCUGUACAGCAU-3’) and its negative control (shNC; 5’-UACGCUCAGCAUGUGUCACUC-3’), miR-34a mimics (5’-UCGUUCGUGAGCACUUGCGACG-3’) and NC mimics (5’-UCGUCGGAUCGACUGAGAUCU-3’), miR-34a inhibitor (5’-AGCCUUGCUGCAGGUGCGCAU-3’) and NC inhibitor (5’-UGCCUUACUGACGGUCGGAGA-3’) were obtained from GenePharm (Shanghai, China). pcDNA3.1 vector (Thermo Fisher) was employed to construct XIST and SMAD2 overexpression vector. The transfection was conducted using Lipofectamine 2000 (Invitrogen).

### RT-qPCR

Total RNAs of anterior lens capsule tissues or SRA01/04 cells were extracted using TRIzol reagent (Invitrogen, USA). The reverse transcription of RNA into cDNA using PrimeScript RT reagent kit (TaKaRa, China). Subsequently, RT-qPCR was performed with SYBR green detection kit (Takara, Japan) and ABI 7500 RT-PCR system (Applied Biosystems, CA, USA). The 2^-ΔΔCt^ method was applied to calculate the relative expression. GAPDH or U6 was detected as the internal references. The following primers were used: XIST forward, 5’-TCAGCCCATCAGTCCAAGATC-3’ and reverse, 5’-CCTAGTTCAGGCCTGCTTTTCAT-3’; miR-34a forward, 5’-ACCCAGTGCGATTTGTCA-3’ and reverse, 5’-ACTGTACTGGAAGATGGACC-3’; SMAD2 forward, 5’-TCCTACTACCGCCTCACA-3’ and reverse, 5’-ACCTCCTCCTCCTCCTCT-3’; GAPDH forward, 5’-TCGACAGTCAGCCGCATCTTCTTT-3’ and reverse, 5’-ACCAAATCCGTTGACTCCGACCTT-3’; U6 forward, 5’-GCTTCGGCAGCACATATACTAAAAT-3’ and reverse, 5’-CGCTTC ACGAATTTGCGTGTCA-3’.

### MTT assay

Cell proliferation was assessed using MTT assay. Transfected SRA01/04 cells were cultured in 96-well plates. MTT (5 mg/ml) was then added for 4h at 37 °C. Next, 150 μL DMSO (Thermo Fisher, USA) was added to each well and the OD490 nm value was measured using a microplate reader (BioTek Instruments, Canada).

### Transwell assay

The invaded SRA01/04 cells were assessed by transwell chambers (8.0 μm pore size; EMD Millipore, USA) and Matrigel (Corning Inc., USA). SRA01/04 cells (2×10^5^ per well) were plated in the upper chamber in serum-free DMEM. 500 μl DMEM (10% FBS) was plated in the lower chambers. After incubation for 48 h at 37°C, invaded cells were stained with 0.1% crystal violet for 20 min and counted under a light microscope (Olympus Corporation).

### Wound healing assay

SRA01/04 cells were plated in 6-well plates and indicated to reach 70% confluence for the wound healing assay. A 200 μL pipette tip was applied to generate artificial scratches. Images of migrated cells were acquired at 0 h and 24 h with the application of a microscope (Leica DMI4000B, Milton Keynes, Bucks, UK).

### TUNEL assay

TUNEL Apoptosis Kit (Roche, Mannheim, Germany) was employed to assess cell apoptosis. After dehydrating by ethanol, SRA01/04 cells were dyed and cultured with TUNEL reaction mixture (Roche). The nuclear staining was performed with DAPI. A microscope (Olympus) was utilized to observe TUNEL-positive cells.

### Dual-luciferase reporter assay

StarBase 2.0 (http://starbase.sysu.edu.cn) was performed to predict the potential bind sites of miR-34a and XIST or SMAD2. The wild-type (wt) or mutant (mut) 3’-UTR sequences of XIST or SMAD2 was cloned into pmirGLO vector (Promega, Madison, WI, USA). The pGL3-XIST-wt or XIST-mut and pmirGLO-SMAD2-wt or SMAD2-mut reporter vector were co-transfected with miR-34a mimics and NC mimics into SRA01/04 cells using Lipofectamine 2000 (Invitrogen). After 48h incubation, the luciferase activity was measured by Dual-Luciferase Reporter Assay System (Promega).

### Statistical analysis

The data were expressed as mean ± SD and performed using SPSS 17.0 (SPSS, Chicago, IL). Each experiment was performed in triplicate. The correlation between miR-34a and XIST or SMAD2 was assessed by Pearson’s correlation analysis. Comparisons between two groups or among multiple groups were assessed using the Student’s t-test or one-way ANOVA. *P*<0.05 was considered significant.

## Results

### XIST is upregulated and miR-34a is downregulated in DC tissues and cells

Firstly, the levels of XIST and miR-34a were detected in DC tissues by RT-qPCR. As exhibited in Figure 1A and B, the level of XIST was elevated and miR-34a expression was reduced in DC samples. Meanwhile, an inverse correlation between the miR-34a level and XIST was discovered in DC tissues (Figure 1C). Moreover, RT-qPCR analysis determined that XIST level was markedly elevated and miR-34a abundance was reduced in SRA01/04 cells treated with HG compared with NG group (Figure 1D and E). The data suggested that XIST and miR-34a might be implicated in DC development.

**Figure 1.**
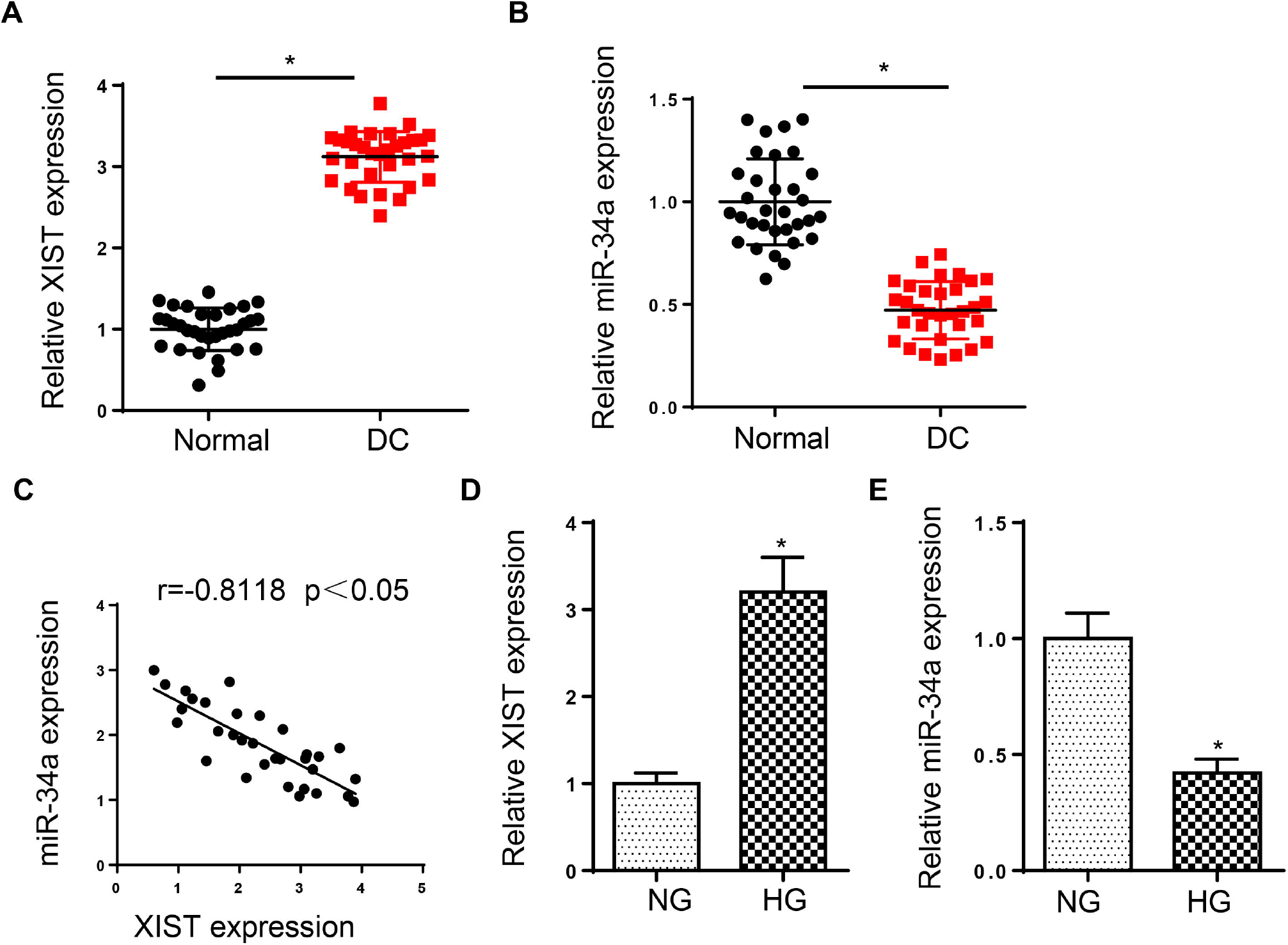
XIST is upregulated and miR-34a is downregulated in DC tissues and cells. (A and B) RT-qPCR showed the relative XIST and miR-34a expression in DC tissues and normal samples. (C) Pearson’s correlation analysis was used to determine the correlation between the miR-34a expression and XIST in DC tissues. (D and E) RT-qPCR showed the relative XIST and miR-34a expression in SRA01/04 cells treated with high glucose. *p<0.05.

### XIST depletion inhibits DC development in HG-induced LECs

To explore the role of XIST in DC, SRA01/04 were transfected with shXIST and shNC before HG treatment. RT-qPCR analysis manifested that XIST depletion decreased the level of XIST in SRA01/04 cells (Figure 2A). Then, MTT and wound healing and transwell assays revealed that XIST deficiency restrained the proliferation, migration and invasion of SRA01/04 cells treated by HG (Figure 2B-D). Meanwhile, XIST depletion induced the apoptosis of HG-stimulated SRA01/04 cells (Figure 2E). These results indicated that XIST regulated HG-treated LECs proliferation, metastasis and apoptosis in DC.

**Figure 2.**
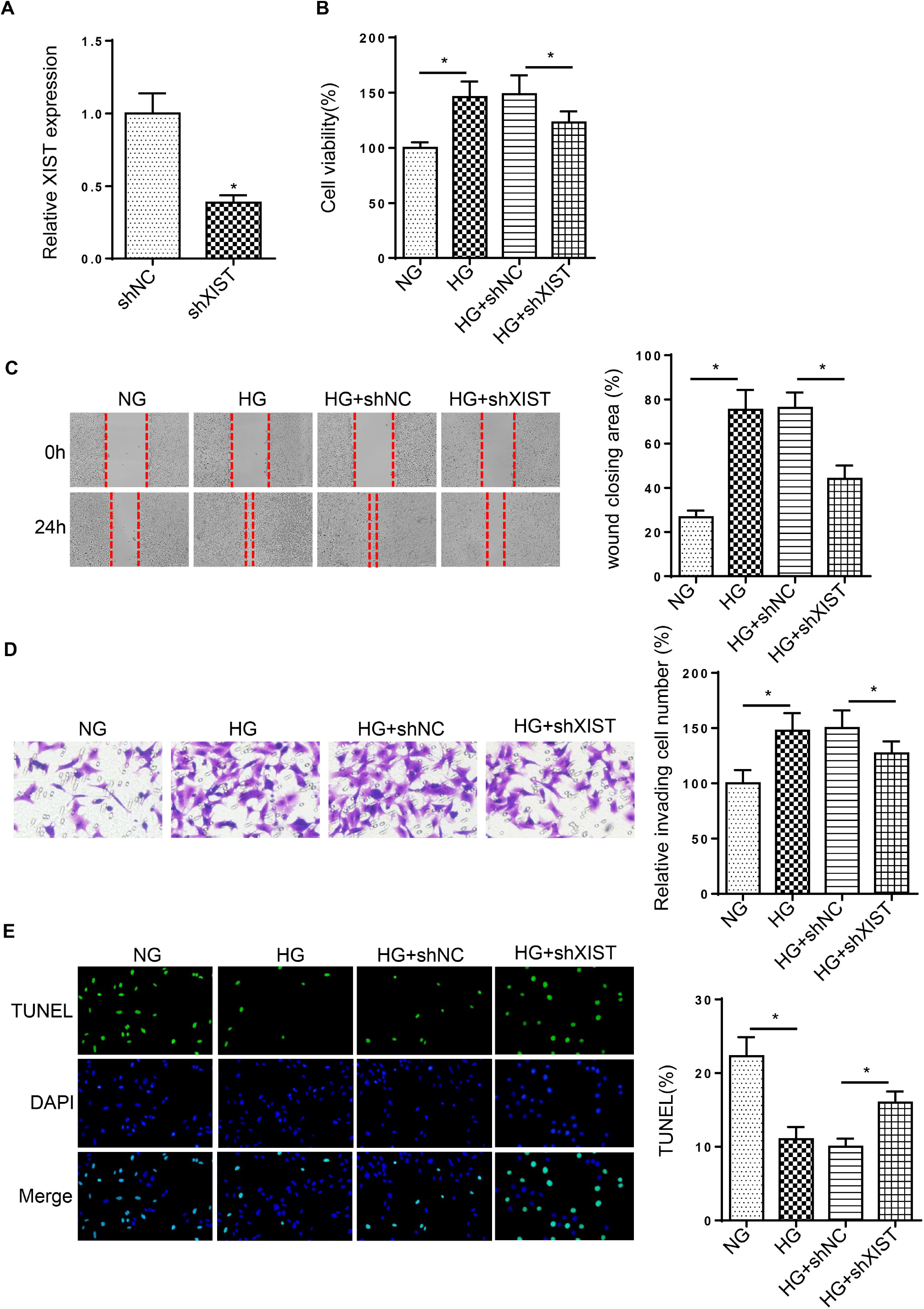
XIST depletion inhibits DC development in HG-induced LECs. (A) RT-qPCR was used to determine the relative XIST expression in SRA01/04 transfected with shXIST. (B-D) MTT and wound healing and transwell assays were used to analyze the proliferation, migration and invasion of SRA01/04 cells transfected with shXIST or shNC after treatment of HG. (E) TUNEL assay was used to determine the apoptosis in SRA01/04 cells transfected with shXIST or shNC after treatment of HG. *p<0.05.

### miR-34a addition weakens HG-mediated the proliferation, migration and invasion in LECs

To investigate the effect of miR-34a on DC, SRA01/04 cells were transfected with miR-34a mimics. As exhibited in Figure 3A, miR-34a was highly expressed by miR-34a addition. Moreover, HG stimulation increased SRA01/04 cell proliferation, migration and invasion, while miR-34a overexpression reversed these effects (Figure 3B-D). Besides, the decreased cell apoptosis caused by HG treatment was rescued by miR-34a addition (Figure 3E). The above discoveries indicated that miR-34a acted as essential roles in HG-treated LECs.

**Figure 3.**
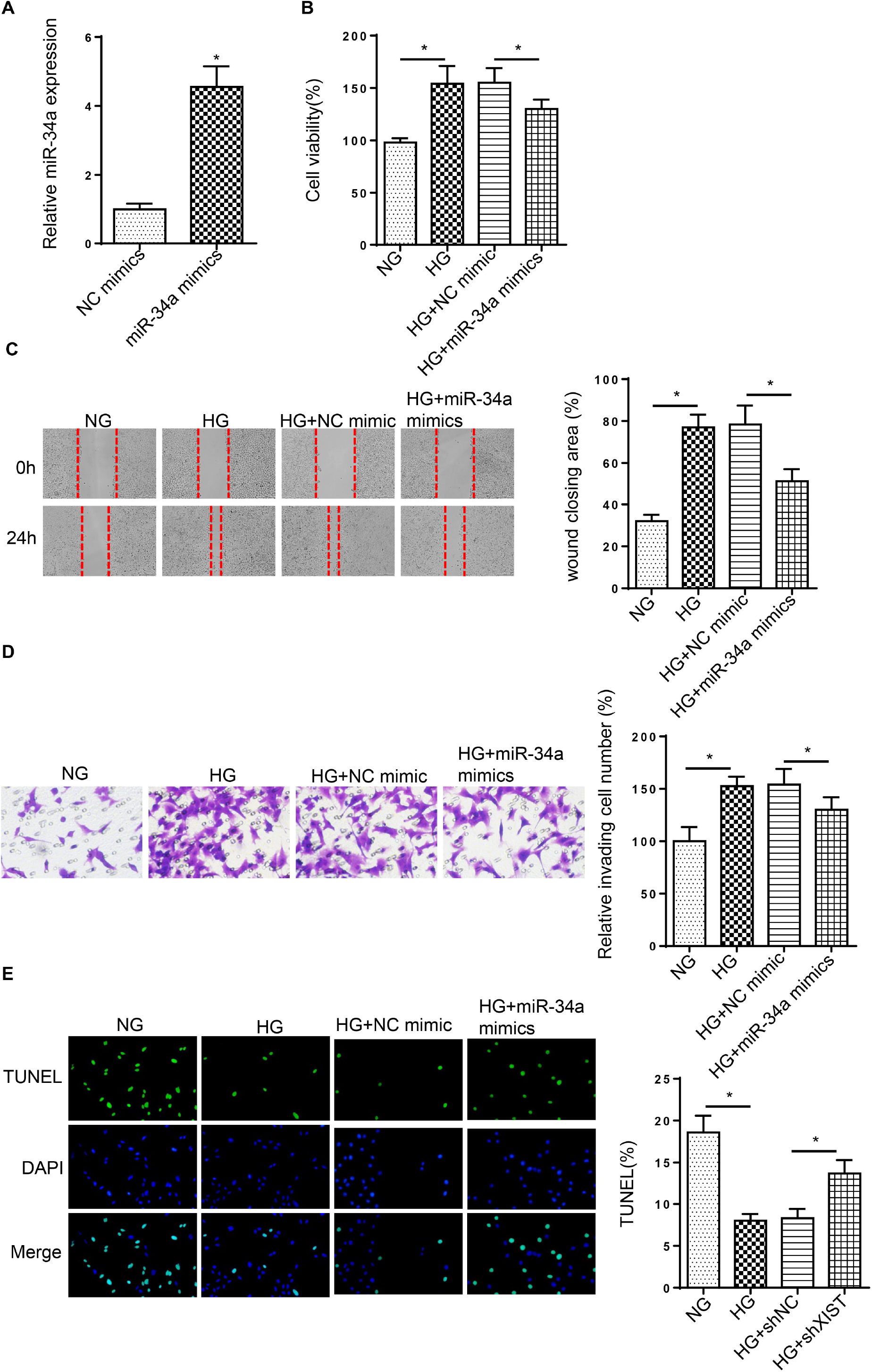
miR-34a addition weakens HG-mediated the proliferation, migration and invasion in LECs. (A) RT-qPCR showed the relative miR-34a expression in SRA01/04 transfected with miR-34a mimics. (B-D) MTT and wound healing and transwell assays were used to analyze the proliferation, migration and invasion of SRA01/04 cells transfected with miR-34a mimics or NC mimics after treatment of HG. (E) TUNEL assay determined the apoptosis in SRA01/04 cells transfected with miR-34a mimics or NC mimics after treatment of HG. *p<0.05.

### XIST deletion modulates DC progression by targeting miR-34a in HG-induced LECs

Subsequently, we explored whether XIST modulated DC progression through miR-34a, As presented in Figure 4A, starBase predicted the binding sequences of XIST and miR-34a. Then, dual-luciferase reporter assay elaborated that miR-34a supplementation repressed the luciferase activity of XIST-wt group, while the activity was unchanged in XIST-mut (Figure 4B). Additionally, RT-qPCR assay demonstrated that the level of miR-34a was reduced by XIST addition in SRA01/04 cells and was enhanced by XIST interference (Figure 4C). Functional assay implied that deficiency of miR-34a rescued the repressive effects of XIST knockdown on the proliferation, migration and invasion in cells stimulated by HG (Figure 4D-F). Meanwhile, XIST silence induced cell apoptosis, which was abrogated by miR-34a deletion in HG-treated SRA01/04 cells (Figure 4G). Therefore, the findings suggested miR-34a was sponged by XIST and participated in the regulation of XIST during DC progression.

**Figure 4.**
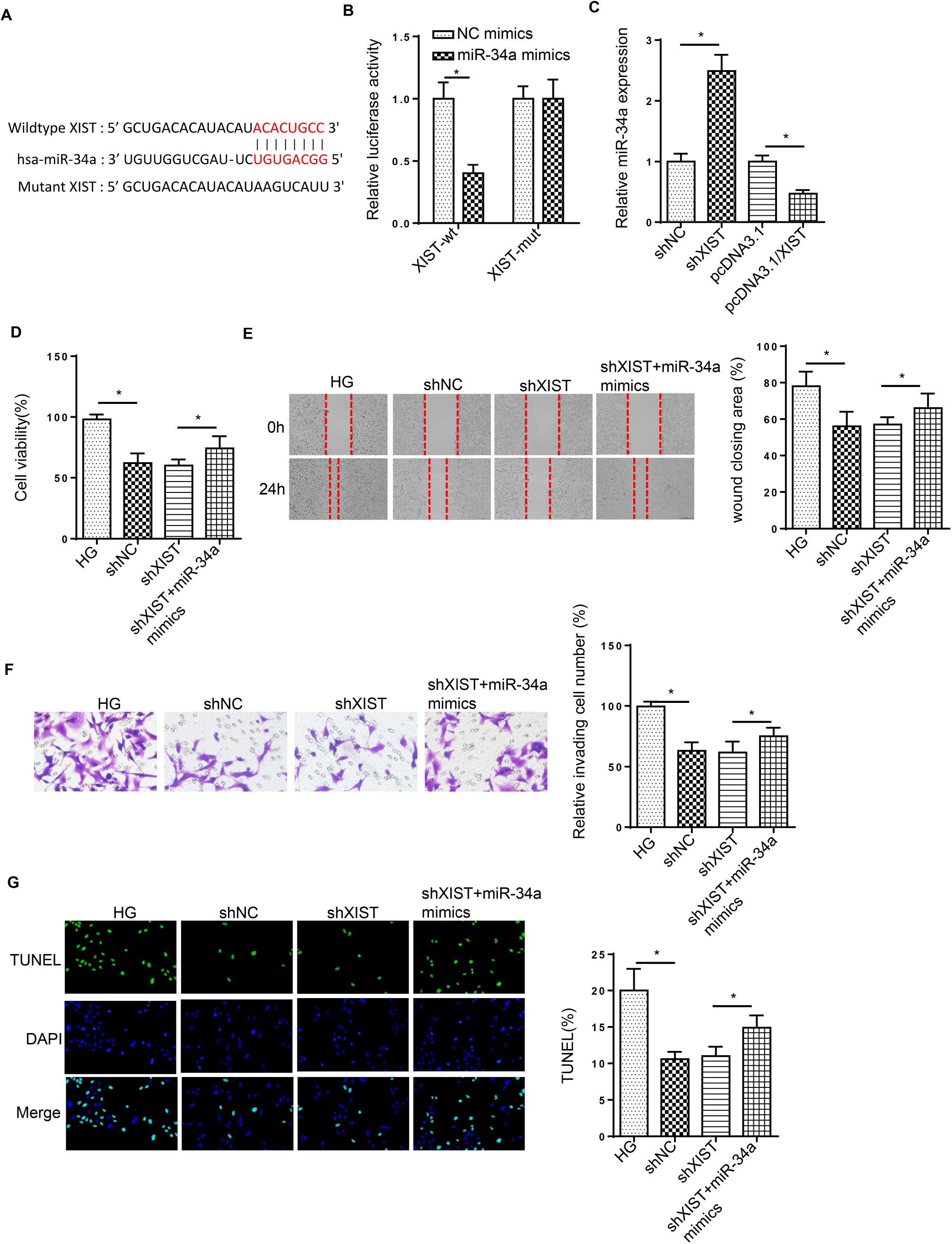
XIST deletion modulates DC progression by targeting miR-34a in HG-induced LECs. (A) StarBase website was used to predict the binding site between XIST and miR-34a. (B) Luciferase reporter assay showed luciferase activity of XIST-wt or XIST-mut in SRA01/04 cells transfected with NC mimics or miR-34a mimics. (C) RT-qPCR showed the relative miR-34a expression in SRA01/04 cells transfection with pcDNA3.1, pcDNA3.1/XIST, shNC or shXIST. (D-F) MTT and wound healing and transwell assays were used to analyze the proliferation, migration and invasion of SRA01/04 cells transfected with shNC, shXIST or shXIST+miR-34a inhibitor after treatment of HG. (G) TUNEL assay determined the apoptosis in SRA01/04 cells transfected with shNC, shXIST or shXIST+miR-34a inhibitor after treatment of HG. *p<0.05.

### SMAD2 is a target of miR-34a

To explore the downstream mechanism of miR-34a in DC progression, starBase website was adopted to predict the putative binding sites of miR-34a and SMAD2 (Figure 5A). Then, the luciferase reporter assay manifested that miR-34a addition restrained the luciferase activity of SMAD2-wt, but not changed the luciferase activity of SMAD2-mut (Figure 5B). Moreover, SMAD2 level in DC tissues was elevated and positively correlated with XIST (Figure 5C and D). Furthermore, higher level of SMAD2 was found in SRA01/04 cells treated by HG than those in NG group (Figure 5E). Besides, SMAD2 expression in SRA01/04 cells was downregulated by miR-34a addition and upregulated by miR-34a deletion (Figure 5F). Above all, miR-34a targeted SMAD2 in SRA01/04 cells.

**Figure 5.**
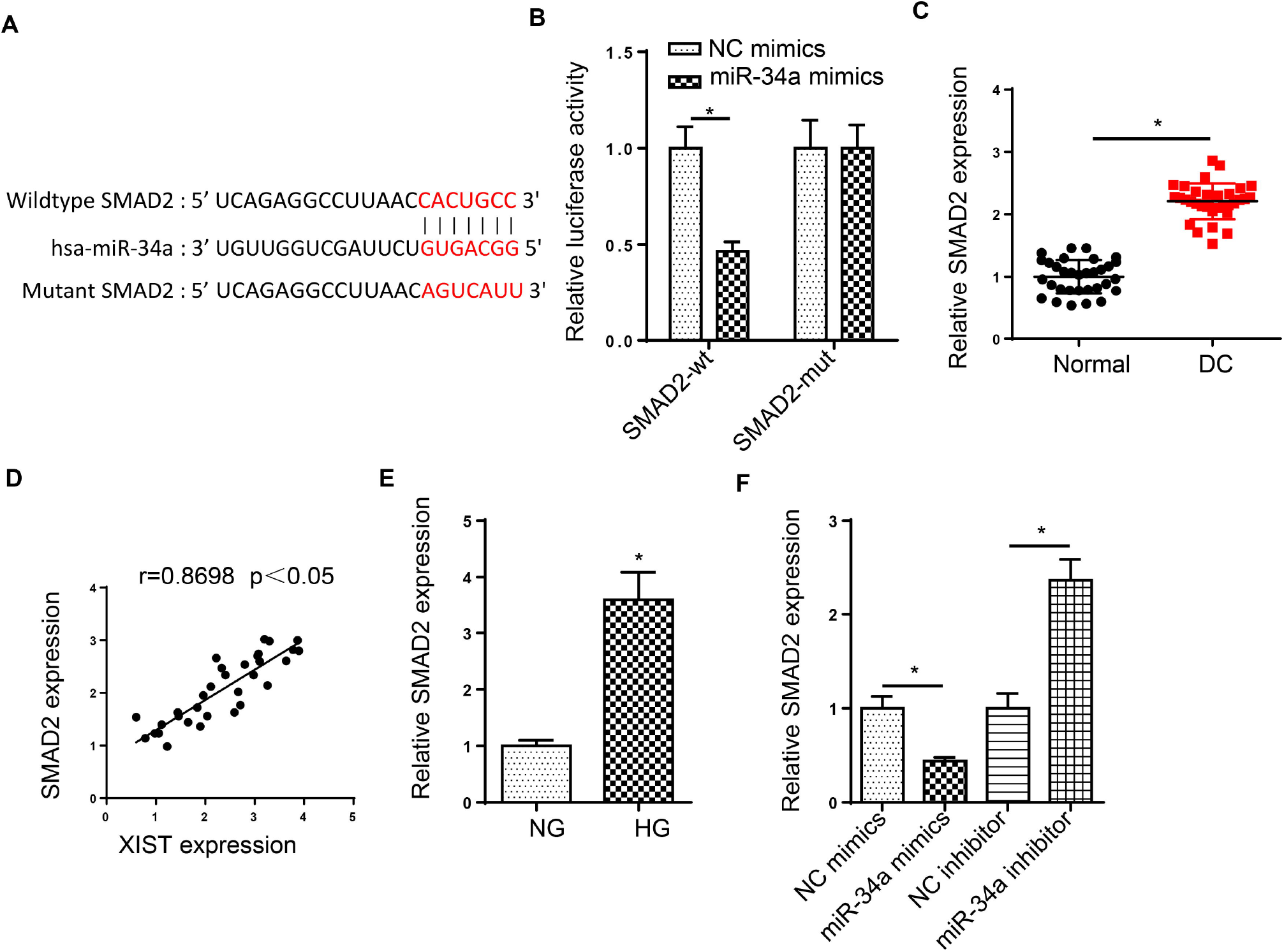
SMAD2 is a target of miR-34a. (A) StarBase website was used to predict the binding site between miR-34a and SMAD2. (B) Luciferase reporter assay showed luciferase activity of SMAD2-wt or SMAD2-mut in SRA01/04 cells transfected with NC mimics or miR-34a mimics. (C and D) RT-qPCR showed the relative SMAD2 expression in DC tissues and normal samples. Pearson’s correlation analysis showed the correlation between the miR-34a expression and SMAD2 in DC tissues. (E) RT-qPCR showed the relative SMAD2 expression in SRA01/04 cells treated with HG. (F) RT-qPCR showed the relative SMAD2 expression in SRA01/04 cells transfected with NC mimics or miR-34a mimics and NC inhibitor or miR-34a inhibitor, *p<0.05.

### XIST accelerates DC progression through SMAD2 by sponging miR-34a in HG-treated LECs

RT-qPCR assay revealed that SMAD2 expression was inhibited by miR-34a supplementation, which was rescued by XIST overexpression in HG-treated cells (Figure 6A). To determine whether XIST mediated DC progression by modulating SMAD2, SRA01/04 cells were transfected with pcDNA3.1, pcDNA3.1/SMAD2 and pcDNA3.1/SMAD2+shXIST. Functional assays elucidated that addition of SMAD2 facilitated cell proliferation, migration and invasion in HG-stimulated SRA01/04 cells, but these effects were abrogated by XIST silence (Figure 6B-D). Furthermore, overexpression of SMAD2 inhibited HG-stimulated SRA01/04 cell apoptosis, which was reversed by XIST silence (Figure 6E). Taken together, XIST contributed to DC development via modulating SMAD2 in HG-induced LECs.

**Figure 6.**
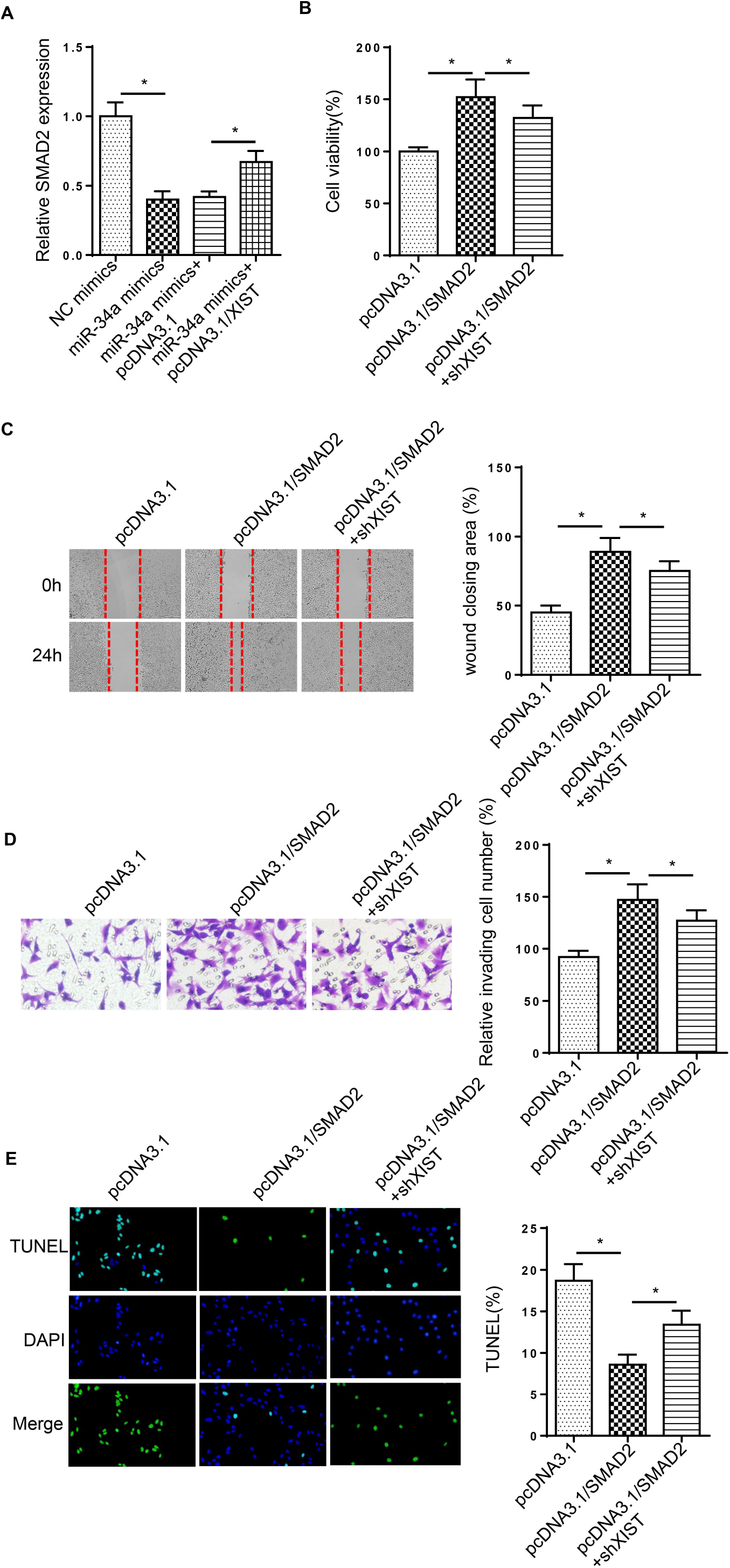
XIST accelerates DC progression through SMAD2 by sponging miR-34a in HG-treated LECs. (A) RT-qPCR analysis determined the SMAD2 expression in SRA01/04 cells transfected with NC mimics, miR-34a mimics, miR-34a mimics+pcDNA3.1/XIST. (B-D) MTT and wound healing and transwell assays were used to analyze the proliferation, migration and invasion of SRA01/04 cells transfected with pcDNA3.1, pcDNA3.1/SMAD2 and pcDNA3.1/SMAD2+shXIST after treatment of HG. (E) TUNEL assay determined the apoptosis in SRA01/04 cells transfected with pcDNA3.1, pcDNA3.1/SMAD2 and pcDNA3.1/SMAD2+shXIST after treatment of HG. *p<0.05.

## Discussion

DC is an early ocular complication in diabetic patients and one of the main causes of blindness [17]. Some literature indicated that lncRNAs affected multiple biological processes in DC [18, 19]. Thus, understanding the function of lncRNA in LECs development under HG is helpful to discover new targets for DC treatment. The current study investigated the role of XIST during DC development. Collectively, the results determined that XIST regulated SMAD2 expression to participate occurrence and development of DC via sponging miR-34a.

The dysregulation of XIST has been found in multiple biological processes, such as proliferation, invasion and apoptosis [20, 21]. XIST was reported to facilitate retinoblastoma progression through sponging miR-101 to mediate the levels of ZEB1 and ZEB2 [22]. Moreover, XIST knockdown repressed retinoblastoma development via the miR-124/STAT3 axis [23]. In the present study, XIST level was enhanced in posterior capsular tissues and HG-stimulated SRA01/04 cells. XIST knockdown attenuated cell proliferation, migration, invasion and induced the apoptosis in LECs induced by HG. These discoveries suggested that XIST and miR-34a might be promising targets for DC treatment.

Extensive evidence has demonstrated that XIST could serve as a ceRNA for miRNAs to regulate various diseases. For example, lncRNA XIST facilitated cell proliferation and invasion via targeting miR-137 to upregulate PXN in non-small cell lung cancer [24]. LncRNA XIST accelerated extracellular matrix degradation by acting as a ceRNA of miR-1277-5p in osteoarthritis [25]. A previous study showed that miR-34a could restrain LEC proliferation and migration by reducing c-Met [26]. In addition, Han et al manifested that miR-34a suppressed EMT of LECs through targeting Notch1 [27]. Herein, miR-34a was downregulated in DC tissues and HG-stimulated SRA01/04 cells, and miR-34a supplementation attenuated DC development in HG-induced LECs. Moreover, deficiency of miR-34a reversed the effects of XIST deletion on cell viability, migration, invasion and apoptosis in cells treated by HG.

Nuclear translocation of SMAD family member 2 (SMAD2) has been found and validated as a crucial regulator associated with cell viability, migration and EMT [28, 29]. The current study confirmed that SMAD2 was the downstream target of miR-34a. Moreover, we also determined that SMAD2 was upregulated DC in tissues and HG-stimulated SRA01/04 cells. Addition of SMAD2 accelerated cell proliferation, migration and invasion and suppressed the apoptosis in SRA01/04 cells under HG treatment. However, these effects were weakened by XIST knockdown. Above all, we illustrated that XIST modulated DC development through regulating SMAD2 in SRA01/04 cells treated by HG.

In summary, this research revealed a novel mechanism of XIST in DC progression. XIST facilitated cell proliferation, migration and invasion as well as decreased apoptosis of HG-induced LECs via miR-34a/SMAD2 axis, revealing a novel biomarker for DC treatment.

## Conflicts of interest

The authors declare that no competing interest exists in this study.

## Funding

None

## Availability of data and materials

The data and materials used in the current study are available from the corresponding author on reasonable request.

## References

1. McCarty, C. A. and H. R. Taylor, Recent developments in vision research: light damage in cataract. Invest Ophthalmol Vis Sci, 1996. 37(9): p. 1720–3.

2. Pollreisz, A. and U. Schmidt-Erfurth, Diabetic cataract-pathogenesis, epidemiology and treatment. J Ophthalmol, 2010. 2010: p. 608751.

3. Harding, J.J., et al., Diabetes, glaucoma, sex, and cataract: analysis of combined data from two case control studies. Br J Ophthalmol, 1993. 77(1): p. 2–6.

4. Kahn, H.A., et al., The Framingham Eye Study, ii. Association of ophthalmic pathology with single variables previously measured in the Framingham Heart Study. Am J Epidemiol, 1977. 106(1): p. 33–41.

5. Martinez, G. and R.U. de longh, The lens epithelium in ocular health and disease. Int J Biochem Cell Biol, 2010. 42(12): p. 1945–63.

6. Wang, Y., et al., Expression Profiling of DNA Methylation and Transcriptional Repression Associated Genes in Lens Epithelium Cells of Age-Related Cataract. Cell Mol Neurobiol, 2017. 37(3): p. 537–543.

7. Mathieu, E.L., et al., [Functions of lncRNA in development and diseases], Med Sci (Paris), 2014. 30(8-9): p. 790–6.

8. Gong, W., et al., LncRNA MALAT1 promotes the apoptosis and oxidative stress of human lens epithelial cells via p38MAPK pathway in diabetic cataract. Diabetes Res Clin Pract, 2018. 144: p. 314–321.

9. Yang, J., S. Zhao, and F. Tian, SP1-mediated lncRNA PVT1 modulates the proliferation and apoptosis of lens epithelial cells in diabetic cataract via miR-214-3p/MMP2 axis. J Cell Mol Med, 2020. 24(1): p. 554–561.

10. Dong, Y., et al., Long non-coding RNA XIST regulates hyperglycemia-associated apoptosis and migration in human retina! pigment epithelial cells. Biomed Pharmacother, 2020. 125: p. 109959.

11. Wang, Q., XIST silencing alleviated inflammation and mesangia! cells proliferation in diabetic nephropathy by sponging miR-485. Arch Physiol Biochem, 2020: p. 1–7.

12. Bartel, D.P., MicroRNAs: genomics, biogenesis, mechanism, and function. Cell, 2004. 116(2): p. 281–97.

13. Zhang, L, R. Cheng, and Y. Huang, MiR-30a inhibits BECN1-mediated autophagy in diabetic cataract. Oncotarget, 2017. 8(44): p. 77360–77368.

14. Zeng, K., et al., Effects of microRNA-211 on proliferation and apoptosis of lens epithelial cells by targeting SIRT1 gene in diabetic cataract mice. Biosci Rep, 2017. 37(4).

15. Xiang, W., et al., miR34a suppresses proliferation and induces apoptosis of human tens epithelial cells by targeting E2F3. Mol Med Rep, 2016. 14(6): p. 5049–5056.

16. Li, H., et al., miR-30a reverses TGF-beta2-induced migration and EMT in posterior capsular opacification by targeting Smad2. Mol Biol Rep, 2019. 46(4): p. 3899–3907.

17. Zhou, Y., et al., Autoregenerative redox nanoparticles as an antioxidant and glycation inhibitor for palliation of diabetic cataracts. Nanoscale, 2019. 11(27): p. 13126–13138.

18. Ye, W., et al., LncRNA MALAT1 Regulates miR-144-3p to Facilitate Epithelial-Mesenchymal Transition of Lens Epithelial Cells via the ROS/NRF2/Notch 1/Snail Pathway. Oxid Med Cell Longev, 2020. 2020: p. 8184314.

19. Li, Y., et al., Roie of incRNA NEAT1 mediated by YY1 in the development of diabetic cataract via targeting the microRNA-205-3p/MMP16 axis. Eur Rev Med Pharmacol Sci, 2020. 24(11): p. 5863–5870.

20. Wang, H., et al., Long non-coding RNA XIST promotes the progression of esophageal squamous cell carcinoma through sponging miR-129-5p and upregu/ating CCND1 expression. Cell Cycle, 2020: p. 1–15.

21. Wang, N., et al., The lncRNA XIST promotes colorectal cancer cell growth through regulating the miR-497-5p/FOXK1 axis. Cancer Cell Int, 2020. 20(1): p. 553.

22. Cheng, Y., et al., LncRNA XIST promotes the epithelial to mesenchymal transition of retinoblastoma via sponging miR-101. Eur J Pharmacol, 2019. 843: p. 210–216.

23. Hu, C., et al., Knockdown of lncRNA XIST inhibits retinoblastoma progression by modulating the miR-124/STAT3 axis. Biomed Pharmacother, 2018. 107: p. 547–554.

24. Jiang, H., et al., Knockdown of long non-coding RNA XIST inhibits cell viability and invasion by regulating miR-137/PXN axis in non-small cell lung cancer. Int J Biol Macromol, 2018. 111: p. 623–631.

25. Wang, T., et al., Long non-coding RNA XIST promotes extracellular matrix degradation by functioning as a competing endogenous RNA of miR-1277-5p in osteoarthritis. Int J Mol Med, 2019. 44(2): p. 630–642.

26. Feng, D., et al., MicroRNA-34a suppresses human lens epithelial cell proliferation and migration via downregulation of c-Met. Clin Chim Acta, 2019. 495: p. 326–330.

27. Han, R., et al., MicroRNA-34a inhibits epithelial-mesenchymal transition of lens epithelial cells by targeting Notch 1. Exp Eye Res, 2019. 185: p. 107684.

28. Li, H., et al., Implication of Smad2 and Smad3 in transforming growth factor-beta-induced posterior capsular opacification of human lens epithelial cells. Curr Eye Res, 2015. 40(4): p. 386–97.

29. Li, J., X. Tang, and X. Chen, Comparative effects of TGF-beta2/Smad2 and TGF-beta2/Smad3 signaling pathways on proliferation, migration, and extracellular matrix production in a human lens cell line. Exp Eye Res, 2011.92(3): p. 173–9.

